# Distinct cortical excitability and connectivity profiles within the human SMA complex

**DOI:** 10.64898/2026.01.30.702764

**Authors:** Francesco Lomi, Ali Jafarov, Fiammetta Iannuzzo, Timo Roine, Ida Granö, Juha Gogulski, Dogu Baran Aydogan, Ilkka Laakso, Mario Rosanova, Simone Rossi, Nikos Makris, Risto J. Ilmoniemi, Pantelis Lioumis

## Abstract

**Introduction and aim:** Understanding brain functions increasingly relies on a network-based perspective, emphasizing interactions across distributed regions. High-order cognitive functions, like language and executive processes, engage such networks rather than isolated cortical areas, yet mapping their dynamics in humans remains challenging. The supplementary motor area (SMA) complex serves as a crucial hub, functioning as an interface between cognitive and sensorimotor processing. Navigated transcranial magnetic stimulation combined with electroencephalography (TMS–EEG) enables causal, time-resolved assessment of cortical excitability and connectivity with high temporal and spatial precision. Extending prior proof-of-concept work, this study aims to provide normative neurophysiological signatures of the SMA complex (pre-SMA and SMA).

**Methods:** Twenty-one healthy subjects underwent a TMS–EEG session where six stimulation targets over the SMA complex (three in pre-SMA, one in the border between pre-SMA and SMA, two in SMA) were recorded. Global cortical excitability via global mean field power (GMFP) was computed over three time-windows (10–50, 50–100, and 100–200 ms) and compared across targets. Normalized power in different frequency bands (α, β1, β2, γ) and natural frequency (NF) were also computed and compared across stimulation sites. Then, we calculated tractography-derived connectivity metrics to examine corticothalamic projections from each stimulation site. Apparent fiber density (AFD)-derived connection strength quantified intra-axonal fibre volume along pathways to the whole thalamus, whereas fibre bundle capacity (FBC) assessed the aggregate intra-axonal cross-sectional area at pathway endpoints, providing an estimate of information transmission capacity to each of the 23 thalamic nuclei.

**Results:** Cortical excitability and oscillatory properties varied significantly across TMS targets. A gradient in cortical excitability was observed, where anterior and middle portions of pre-SMA exhibited significantly lower GMFP in the early post-stimulus phase (i.e., 10–50 ms) than SMA portions; interestingly, these differences were clearly dampened in a later time-window (50–100 ms) and disappeared after this period. Furthermore, higher α and lower β2 power were observed in anterior pre-SMA than posterior pre-SMA portions. No differences were observed in NF. Crucially, we also observed significant differences in whole- and single-nuclei thalamic structural connectivity.

**Discussion:** These findings provide direct human *in vivo* evidence for distinct neurophysiological and structural connectivity profiles of the pre-SMA and SMA, possibly reflecting their different cytoarchitectonic, myeloarchitectonic and connectivity properties. Variations in cortical excitability, oscillatory dynamics, and FBC along the rostro-caudal axis may mirror functional specialization, with pre-SMA supporting cognitive control processes and SMA engaging motor-related thalamocortical networks. Together, the results demonstrate the sensitivity of TMS–EEG and diffusion-weighted connectomics to measure fine-grained neurophysiological differences of corticothalamic projections, which can lead to individualized neuromodulation targeting.

## 1. INTRODUCTION

Over the past decades, neuroscience has progressively shifted from the single cortical area perspective, in which specific brain areas are thought to underlie distinct cognitive functions, toward a network-based view of brain organization (Bullmore & Sporns, 2012). In this framework, neural functions are understood as emergent properties of large-scale networks composed of spatially distributed but functionally interconnected regions. When one or more hubs of these networks become dysfunctional, a combination of symptoms may manifest a specific psychiatric or neurological disorder (Vogel et al. 2023).

Studying brain networks requires an understanding of both structural and functional architectures. Structural connectivity refers to the anatomical connections linking different brain regions, typically examined using magnetic resonance imaging (MRI) techniques such as diffusion-weighted imaging (DWI). Functional connectivity, by contrast, captures the statistical dependencies between spatially distinct regions during rest or task performance, and can be explored through functional MRI (fMRI) or electrophysiological methods. Recent advancements in neuroimaging have brought renewed attention to this principle (Glasser et al. 2016; Van Essen et al., 2019). Yet, in the human brain, precise knowledge of structural connections from origins to terminations, is still limited (Rushmore et al., 2020), resulting in a partial and not definitive mapping of human brain connectivity.

Among the available techniques, the combination of transcranial magnetic stimulation (TMS) with electroencephalography (EEG) provides a unique opportunity to probe brain connectivity in an active and causal manner (Ilmoniemi et al., 1997; Massimini et al., 2005). By applying focal perturbations and recording the resulting time-resolved cortical responses (Rosanova et al., 2009), TMS–EEG allows to infer the directionality and dynamics of interactions between brain regions with millisecond temporal precision. Recent findings also indicate that TMS–EEG provides a sensitive tool for probing brain activity with high spatial resolution (Passera et al., 2022; Lioumis et al., 2025), reinforcing its potential for mapping functional brain networks beyond the motor system. TMS–EEG could hence complement MRI-based measures by offering a direct and spatiotemporally fine-grained assessment of local cortical excitability and effective connectivity.

Despite single-pulse TMS directly exciting only neocortical structures, subcortical regions can also be engaged through network-mediated effects. In this context, the thalamus plays a crucial role in regulating and coordinating large-scale brain activity (Shine et al., 2023). Converging evidence indicates that TMS-induced EEG responses are not confined to cortico-cortical interactions, but critically involve thalamo-cortical circuits (Claar et al., 2023; Russo et al., 2025). Thalamic structures both receive inputs from the cortex (mainly layer V and VI) and project back to the cortex, thereby forming cortico-thalamo-cortical loops that are essential for communication across hierarchically organized cerebral networks (Sheperd & Yamawaki, 2021). The thalamus comprises approximately 30 distinct nuclei with varying degrees of functional specialization, including first-order and higher-order nuclei, which differentially contribute to the propagation and modulation of cortical activity (Marcuse et al., 2025).

These considerations are central to the concept of target engagement in neuromodulation. Since neuromodulation operates through neuroplastic mechanisms akin to Hebbian learning, target engagement should not be defined solely by the cortical area directly stimulated. Rather, it must also encompass the broader network of brain circuits functionally and structurally connected to that area. On a single-subject basis, this approach considers individual differences in brain anatomy and function that may critically affect the individual response to neuromodulation therapies. However, this idea is still a matter of debate due to methodological challenges (Cocchi & Zalesky, 2018; Soleimani et al., 2025), and standard stimulation protocols based on anatomical landmarks are still commonly adopted.

The present study aimed to map a specific portion of the superior frontal gyrus encompassing the pre-SMA and SMA with TMS–EEG to assess cortical excitability differences across adjacent stimulation targets. The additional contribution of DWI allowed testing whether the observed differences in the EEG responses between targets were explained by their different structural connectivity profiles. We focused specifically on the thalamus given its known role in mediating the observed EEG responses (Claar et al., 2023; Llinàs et al., 2007; Russo et al., 2025).

## 2. METHODS

### 2.1 Participants

Twenty-one subjects (31±7 years old, 9 females) completed the experiment. Inclusion criteria were the following: a) age between 18 and 65 years; b) no neurological or psychiatric conditions; c) structural MRI scan of the subject at least with T1 and DWI sequences available; d) right-handedness; e) no ongoing medications affecting the central nervous system; f) no contraindication to TMS. All participants provided written informed consent before study inclusion, and the experiments were approved by the Coordinating Ethics Committee of the Hospital District of Helsinki and Uusimaa (HUS/1198/2016).

### 2.2 MRI acquisition

MRI data were acquired at the Advanced Magnetic Imaging Centre (Aalto University, Espoo, Finland) using a 3-tesla Siemens Skyra scanner (Siemens Healthineers, Erlangen, Germany). Participants’ heads were stabilized with foam padding to minimize motion artifacts, and they were instructed to remain awake and relaxed throughout the scanning session.

High-resolution T1-weighted anatomical images were obtained using a three-dimensional magnetization-prepared rapid gradient-echo sequence with an isotropic voxel resolution of 1 mm. T2-weighted images were acquired with a three-dimensional turbo spin-echo sequence. Multi-shell DWI was performed using an echo-planar imaging sequence with 2 mm slice thickness, comprising 116 diffusion-encoding directions distributed across three b-value shells (900, 1,600, and 2,500 s/mm²) and 10 non-diffusion-weighted (b = 0 s/mm²) reference volumes.

### 2.3 TMS–EEG procedure

Before the TMS–EEG session, the pre-SMA and SMA regions were manually delineated on each participant’s T1-weighted anatomical image following established anatomical landmarks, as described in our previous work (Lioumis et al. 2025). These anatomical masks were then loaded into the neuronavigation system to ensure consistent targeting.

TMS pulses were delivered with a 70-mm radius figure-of-eight coil attached to a Nexstim TMS stimulator (NBT 2.2.4, Nexstim Plc, Helsinki, Finland) with an integrated MRI-guided neuronavigation and electromyography device. The TMS-induced electric field (E-field) strength was monitored in real time using a spherical head model (Hannula & Ilmoniemi, 2017). EEG was recorded using a TMS-compatible amplifier (Bittium Plc, Finland) equipped with a 64-channel cap in a standard montage with passive C-ring-shaped electrodes. The signal was lowpass-filtered at 1250 Hz and sampled at 5000 Hz. Reference and ground electrodes were located on the right mastoid and zygomatic bone, respectively. The skin under each electrode was prepared with conductive abrasive paste (OneStep AbrasivPlus, H + H Medical Devices, Germany) before being filled with conductive gel (Electro-Gel, ECI, Netherlands). The impedances at all electrodes were kept below 5kΩ throughout the experiment. TMS-evoked auditory artifacts were minimized using a TMS-specific noise delivered with a dedicated toolbox (TAAC) (Russo et al., 2022), and played with noise-attenuating in-ear headphones (ER3C Insert Earphones, Etymotic Research Inc., United States). The quality of the TMS-evoked EEG response was ensured by employing a real-time graphical interface displaying the signal in average reference (Casarotto et al., 2022; Parmigiani et al., 2025; Ukharova et al., 2025). The individual stimulation parameters were then adjusted in real time based on the average responses to 20 stimuli.

Prior to initiating TMS–EEG mapping, the peeling depth in the neuronavigation system was set between 20 and 25 mm to ensure clear visualization of the SMA and pre-SMA. Initial stimulus intensity was set approximately at the individual resting motor threshold for the right abductor pollicis brevis muscle, previously identified by means of a dedicated algorithm embedded in the neuronavigation system (Awiszus, 2003). The coil was initially placed orthogonally to the midline in light of evidence showing that the TMS-evoked response is maximized with this orientation (Tervo et al., 2022). The optimal stimulation intensity was then slightly adjusted until a peak-to-peak of at least 10 microvolts was obtained in the channels under the coil (e.g., F1, FC1) for a cortical site in either SMA or preSMA, while minimizing artifacts. This target served as a reference for the electric field (E-field) strength that was kept the same for all other targets. Coil orientation was also modified in case of the occurrence of evoked muscle or decay artifacts. When the above peak-to-peak criterion was reached, we manually set six targets inside the area of interest (Figure 1A): anterior pre-SMA (pre-SMAa), middle pre-SMA (pre-SMAm), posterior pre-SMA (pre-SMAp), pre-SMA/SMA border, anterior SMA (SMAa), posterior SMA (SMAp). In all subjects, targets were identified in left SMA and pre-SMA, except for one participant where targets were moved in the right homologous region due to suboptimal EEG signal quality. Prior to recording, we delivered a few pulses for each target to check whether the artifact levels were minimal; if not, we slightly adjusted the target position. 100 pulses were delivered to each target while keeping the coil position and orientation, the stimulation depth and the electric field (E-field) strength at the target constant. When needed, the stimulation intensity was slightly adjusted before recording to achieve identical E-field strength across targets.

**Figure 1.**
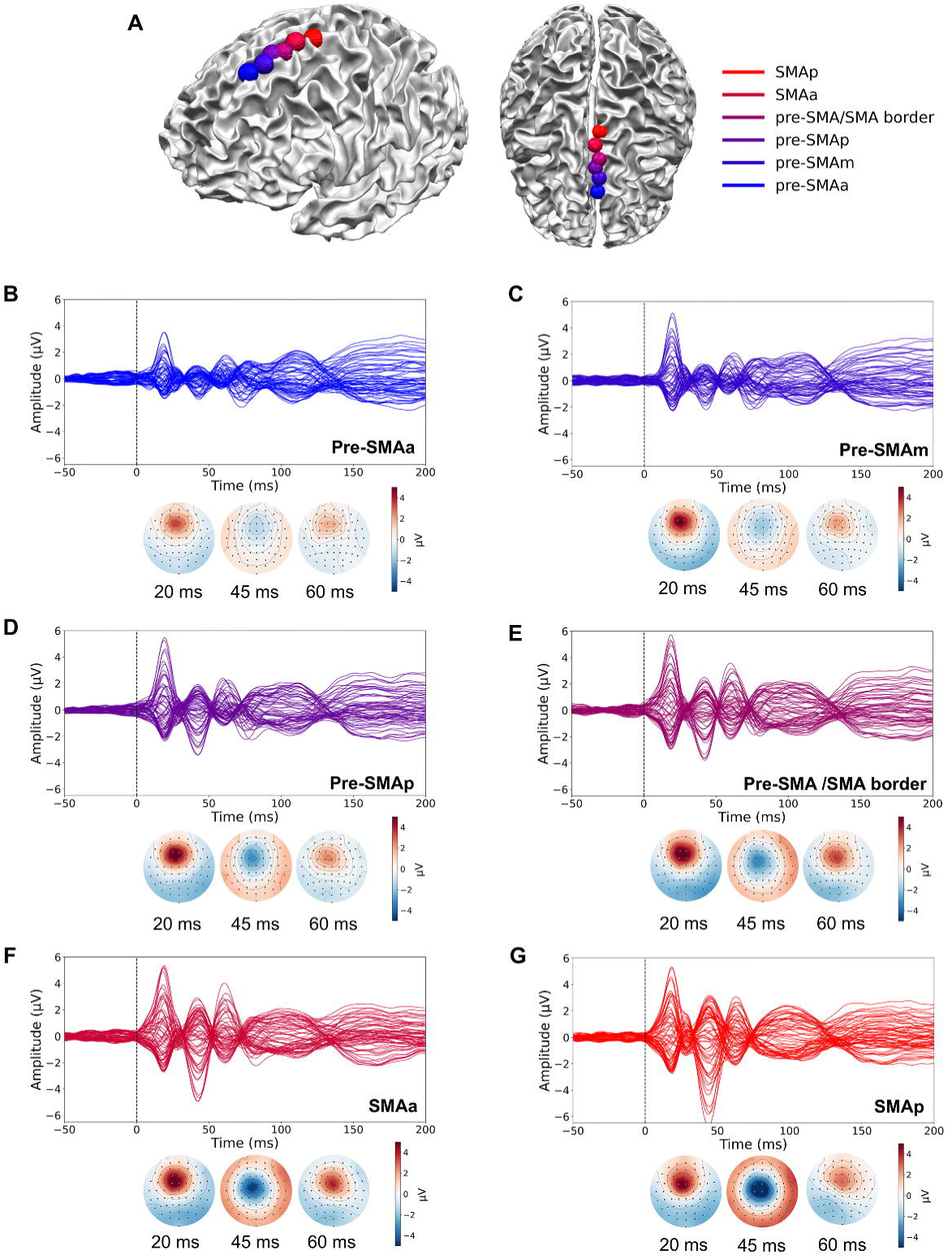
Grand-average TMS–evoked EEG responses across stimulation targets. A) Stimulation locations shown on white matter brain surface in two different views. Panels **B-G** show grand-average butterfly plots of EEG signals time-locked to the TMS pulse for each stimulation target, averaged across subjects. For each target, the corresponding grand-average scalp topographies at 20 ms, 45 ms, and 60 ms post-stimulation are displayed alongside the butterfly plots, illustrating the spatial distribution of TMS-evoked cortical activity across the scalp. Target color coding from blue to red indicates anterior–posterior target position (blue = most anterior, red = most posterior). Pre-SMAa = anterior pre-SMA, pre-SMAm = middle pre-SMA, pre-SMAp = posterior pre-SMA, SMAa = anterior SMA, SMAp = posterior SMA.

### 2.4 TMS–EEG data preprocessing

EEG data were preprocessed in MATLAB R2021b using a custom script based on EEGLAB (v2024.2) (Delorme & Makeig, 2004) and the TESA toolbox together with the FastICA package (Rogasch et al., 2017). Continuous EEG was epoched from −1500 to +1500 ms around the TMS pulse. Each epoch was baseline-corrected using the prestimulus interval from −500 to −5 ms relative to the TMS trigger. The TMS pulse artifact (−3 to 10 ms) was removed and the missing data segment interpolated using cubic spline interpolation. Bad channels (e.g., with big decay artifacts or signal drifts) and noisy trials were identified by visual inspection and removed. To remove ocular artifacts and TMS-independent muscle activity, data dimensionality was reduced via principal component analysis (PCA; 30 components). and FastICA was applied. Artifactual components were identified and removed based on temporal, spectral, and spatial features. When a TMS-evoked muscle artifact was present, the Signal-Space Projection Source-Informed Reconstruction (SSP–SIR) method was applied (Mutanen et al., 2016). Data were subsequently band-pass filtered between 1 and 100 Hz using EEGLAB’s FIR filter, with an additional 50 Hz notch filter to attenuate line noise. Then a second removal of the TMS artifact interval (−3 to 12 ms) was performed, with cubic spline interpolation of the missing data, to eliminate residual TMS-evoked and high-frequency muscle artifacts that were not fully removed in the first step. Data was then resampled to 1000 Hz and re-epoched to −1000 to +1000 ms to avoid edge artifacts from filtering, and average re-referenced. Bad channels were finally interpolated using the original electrode montage with a spherical spline method.

### 2.5 TMS–EEG data analysis

Global mean field power (GMFP) was computed as the root-mean-square deviation of each electrode’s voltage from the instantaneous spatial mean across the scalp (Lehmann & Skrandies, 1980). Mean GMFP values were extracted in three post-stimulus windows (10–50, 50–100, and 100–200 ms). Local mean field power (LMFP) values were also extracted for the same time windows over a region of interest (ROI) covering the area under the coil (i.e., F1, Fz, FC1, FCz, C1, Cz). The channel with the highest peak-to-peak amplitude was selected for each time window within this ROI for each subject and compared between targets.

TMS-evoked peak-to-peak amplitudes were compared between targets over the channel with the highest peak-to-peak in the same time window.

Normalized evoked power was also computed for the same ROI by using the EEGLAB (v2024.2) *newtimef* function on the cleaned data. A Morlet wavelet transform was applied using frequencies ranging from 8 to 45 Hz (1 Hz resolution) and a variable cycle length (3 cycles at the lowest frequency, increasing by 0.5 cycles per frequency increment). The baseline interval was set from −500 to −100 ms. Mean event-related spectral perturbations (ERSPs) were calculated in the 20–200 ms post-stimulus window across predefined frequency bands and averaged across ROI channels: alpha (8–12 Hz), low beta (13–20 Hz), high beta (21–30 Hz), gamma (31–45 Hz) (Rosanova et al., 2009). For each subject, band-specific power values were normalized by expressing them relative to the mean broadband power. The frequency showing the largest cumulated ERSP was defined as the natural frequency (Rosanova et al., 2009; Ferrarelli et al., 2012).

### 2.5 MRI data processing

#### 2.5.1 Anatomical image processing

Cortical surface reconstruction and subcortical volumetric segmentation were performed using FreeSurfer image analysis software (v7.4.1, Fischl B. 2012, open-source at http://surfer.nmr.mgh.harvard.edu). Subcortical segmentations were generated for thalamic nuclei (Iglesias et al., 2018), amygdalar nuclei, and hippocampal subfields (Saygin et al., 2017) using specialized subfield segmentation modules. Thalamic nuclei segmentation yielded 46 nuclei in total (23 left, 23 right), and downstream analyses were conducted at the level of those segmented nuclei.

#### 2.5.2 DWI processing

DWIs were preprocessed using MRtrix3 (v3.0.8) (Tournier et al., 2019). Thermal noise was removed using Marchenko-Pastur PCA-based denoising (Veraart et al., 2016; Cordero-Grande et al., 2019), followed by correction for Gibbs ringing artefacts (Kellner et al., 2016). Geometric distortions arising from susceptibility-induced field inhomogeneities, eddy currents, and subject motion were corrected using the MRtrix3 DWI preprocessing pipeline (Smith et al., 2004; Andersson & Sotiropoulos, 2015), which interfaces with FSL (the FMRIB Software Library; Jenkinson et al., 2012). B1-field inhomogeneity correction was subsequently applied using the N4 bias-field-correction algorithm from Advanced Normalization Tools (Tustison et al., 2010).

Response functions for spherical deconvolution were estimated using the Dhollander algorithm, which performs unsupervised estimation of white matter, grey matter, and cerebrospinal fluid response functions from multi-shell data (Dhollander et al., 2016). To enable group-level comparisons and maintain consistency across subjects, a group-average response function was computed from the individual subject estimates (Smith et al., 2022). Fiber orientation distributions (FODs) were then estimated for each subject using multi-shell multi-tissue constrained spherical deconvolution, which models multiple tissue types simultaneously to improve FOD estimation accuracy in regions with partial volume effects (Tournier et al., 2004, Jeurissen et al., 2014).

Anatomically-Constrained Tractography (ACT; Smith et al., 2012) was implemented to improve the biological plausibility of streamline reconstructions. A five-tissue-type (5TT) segmentation image was generated using segmentation results from FreeSurfer to create tissue probability maps for cortical grey matter, subcortical grey matter, white matter, and cerebrospinal fluid. The Hybrid Surface and Volume Segmentation (Smith et al., 2020; Ardekani et al., 2009; Zhang et al., 2001) algorithm was employed with enhanced segmentation of hippocampal subfields and thalamic nuclei to improve anatomical precision.

Whole-brain probabilistic tractography was performed using second-order integration over fiber orientation distributions (Tournier et al., 2010; Hagmann et al. 2009). Streamline propagation was constrained using the ACT framework with the previously generated 5TT segmentation, which enabled anatomically informed termination and backtracking when streamlines entered inappropriate tissue types. Tractography parameters were set to generate 50 million streamlines with a step size of 0.5 mm, maximum curvature angle of 45° between successive steps, and an FOD amplitude cutoff of 0.06. Spherical-deconvolution-informed filtering of tractograms was applied to assign quantitative weights to each streamline, accounting for the reconstruction biases inherent in probabilistic tractography (Smith et al., 2015).

#### 2.5.3 Quantification of corticothalamic connectivity

ROIs corresponding to six stimulation targets were generated using a custom MATLAB script (R2023a). Spherical ROIs with a 6-mm radius were constructed, centered at coordinates corresponding to the maximum E-field intensity locations (Figure 4A). Only grey-matter voxels within each spherical mask were retained and saved as final ROI masks.

Structural connectivity between cortical ROIs and thalamic structures was extracted from the whole-brain tractogram by selecting streamlines with both termination points residing within the respective ROI pairs. Only direct connections between cortical targets and thalamic nuclei were considered. Then, we extracted the streamlines’ endpoints as a mask and applied Gaussian smoothing at 1mm full-width-at-half-maximum. After transferring the following masks onto the brain surfaces, we considered them as structural connectivity projections (Figure 4B). Voxels with smoothed endpoint density values below 0.3 streamlines were excluded to focus on regions with stronger anatomical projections.

For each of the six targets, average structural connectomes were generated by first computing individual connectivity matrices from extracted tract bundles between each stimulation site and the left segmented thalamus for all except one subjects (□_□_= 20), then averaging these matrices across participants. One participant was excluded from the tractography analysis because the corresponding stimulations were delivered to the right-hemispheric homologues of the predefined target regions. Connectivity was quantified using the sum of per-streamline values for all streamlines assigned to each edge. The resulting weighted connectivity between each stimulated site and the 23 left thalamic nuclei was normalized to the local maximum and visualized using connectograms. Connectivity strengths were then normalized to the maximum value observed across all targets to enable direct comparison between stimulation sites, highlighting both the alterations in connection patterns between stimulation sites and changes in connectivity magnitude.

Apparent Fibre Density (AFD)-derived connection strength was quantified to assess the intra-axonal fibre volume occupied by white matter pathways connecting six spatially neighboring cortical regions to the thalamus. Fibre Bundle Capacity (FBC) was computed to estimate the information transmission capacity of white matter pathways connecting stimulation targets to individual thalamic nuclei.

To exclude that connectivity differences between targets with the thalamus could reflect nonspecific global connectivity differences, total streamlines count for the whole left hemisphere was extracted for each target.

### 2.6 Statistical analysis

Given the violation of the normality assumption in several conditions (Shapiro–Wilk test, *p* < 0.05), non-parametric tests were adopted throughout. Specifically, Friedman tests were used to evaluate within-subject differences in TMS–EEG and DWI (AFD, FBC, streamline count) measures across targets, followed by Wilcoxon signed-rank post-hoc tests with Bonferroni correction. Effect sizes were estimated using Kendall’s *W*.

For each thalamic nucleus, within-subject differences in FBC across stimulation targets were assessed using Friedman tests, with *p*-values corrected for multiple comparisons across nuclei using the Bonferroni correction. Significant Friedman tests were followed by pairwise Wilcoxon signed-rank tests between stimulation targets, with another Bonferroni correction applied to control for multiple post-hoc comparisons.

To investigate the relationship between GMFP in post-TMS time windows (10–50, 50–100, and 100–200 ms) and the AFD-derived connectivity strength of the tract connecting the stimulated cortical area to the thalamus, a linear mixed-effects model (LMM) was employed, including random intercepts for subjects and for targets nested within subjects. Initial residual diagnostics indicated heteroscedasticity, so a generalized least squares (GLS) model with an exponential variance structure was used to account for non-constant residual variance. Fixed effects included AFD-derived connectivity, time-window, and their interaction. The slope of GMFP as a function of AFD-derived connectivity for each time window was estimated using the *emtrends* function from the *emmeans* R package.

## 3. RESULTS

Butterfly plots and scalp topographies for each target are reported in Figure 1. Maximum E-field values were stable across targets (91.61–92.2 V/m range), achieved with stimulation intensities ranging between 56 and 60% of the maximum stimulator’s output (MSO) (i.e., 110–120% range of resting motor threshold, which was at 50.25±8.00 of MSO). Additional information about TMS parameters is reported in supplementary materials (Table S1).

### 3.1 Cortical excitability

Our results (Figure 2–3) demonstrate significant differences of the TMS-evoked response across neighboring cortical locations in the SMA complex (Fig 1A). In particular, TMS-evoked global cortical reactivity at early latencies (i.e., responses within the first 10–50 ms) was stronger when posterior targets were stimulated. These differences became weaker for the later responses across all the targets.

**Figure 2.**
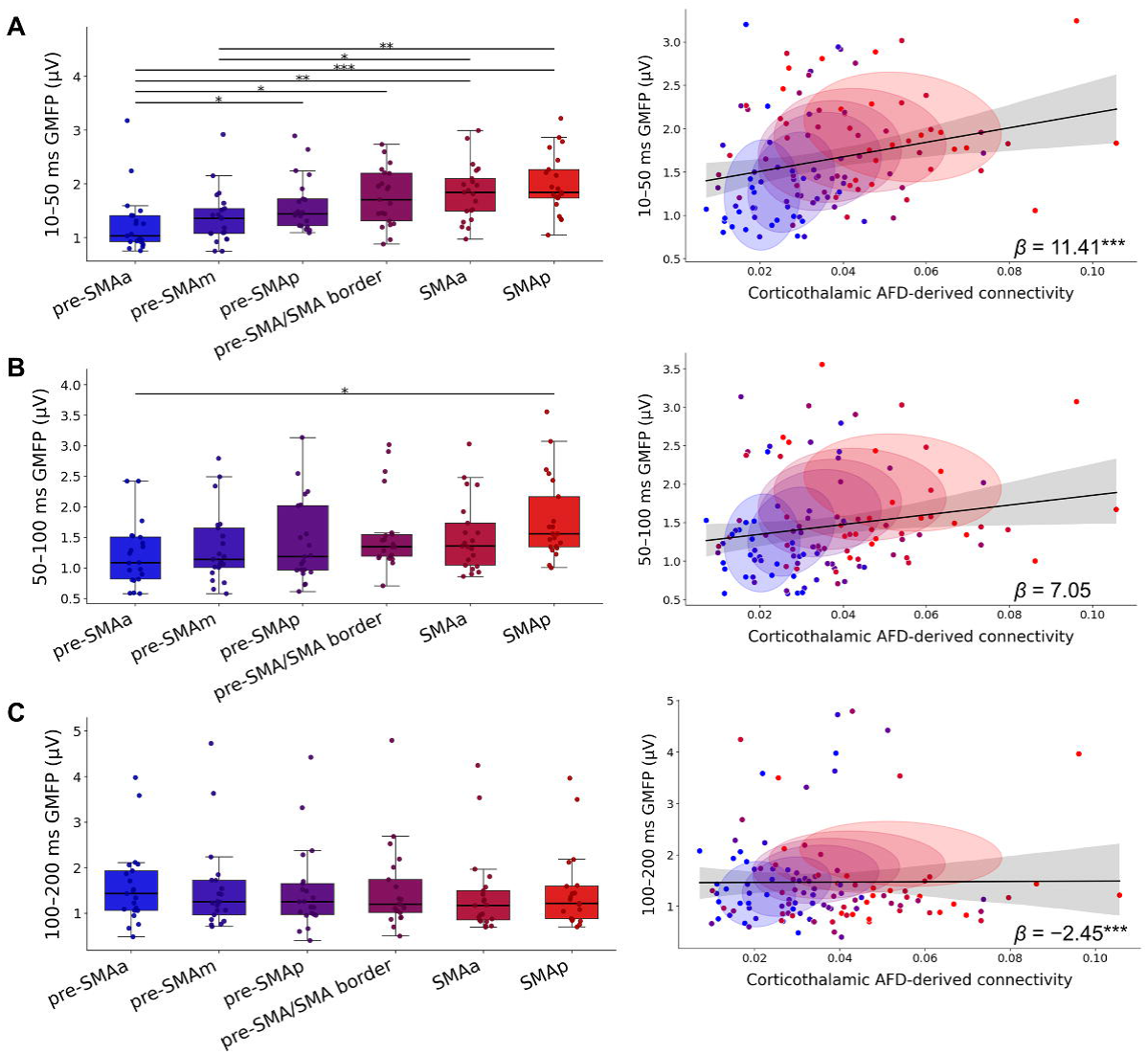
GMFP and its relationship with corticothalamic AFD-derived connectivity across stimulation targets. Panels **A–C** show box plots of GMFP for each target (left) and scatter plots of AFD-derived connectivity versus GMFP with regression lines (right). **A**) GMFP calculated between 10–50 ms post-TMS; **B**) GMFP calculated between 50–100 ms; **C**) GMFP calculated between 100–200 ms. Asterisks indicate significant post-hoc comparisons (Bonferroni-corrected): \*\*\**p* < 0.001, \*\**p* < 0.01, \**p* < 0.05. Target color coding from blue to red indicates anterior–posterior target position (blue = most anterior, red = most posterior). Pre-SMAa = anterior pre-SMA, pre-SMAm = middle pre-SMA, pre-SMAp = posterior pre-SMA, SMAa = anterior SMA, SMAp = posterior SMA.

**Figure 3.**
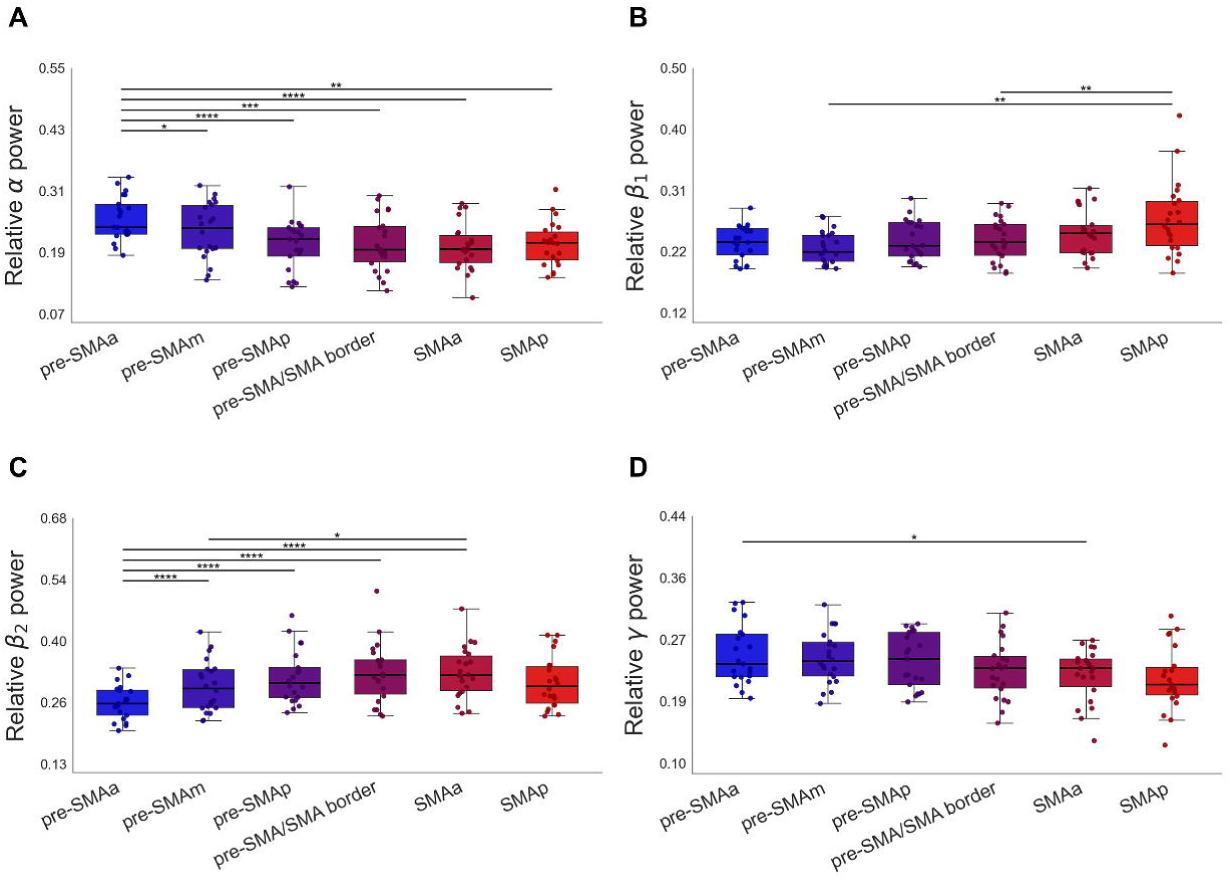
Spectral power across frequency bands. Box plots show the spectral power for each stimulation target in four frequency bands: α, β1, β2, and γ. Asterisks indicate significant post-hoc comparisons (Bonferroni-corrected): \*\*\**p* < 0.001, \*\**p* < 0.01, \**p* < 0.05. Target color coding from blue to red indicates anterior–posterior target position (blue = most anterior, red = most posterior). Pre-SMAa = anterior pre-SMA, pre-SMAm = middle pre-SMA, pre-SMAp = posterior pre-SMA, SMAa = anterior SMA, SMAp = posterior SMA.

More specifically, in the 10–50 ms time window, GMFP was modulated by target location (χ*²*(5) = 54.5, *p* < 0.001) (Figure 2A). The effect size was large (Kendall’s *W* = 0.519) and GMFP was significantly different between pre-SMAa and pre-SMAp (*p* = 0.032), pre-SMAa and pre-SMA/SMA border (*p* = 0.015), pre-SMAa and SMAa (*p* = 0.002), and pre-SMAa and SMAp (*p* < 0.001). In addition, differences were found between pre-SMAm and SMAa (*p* = 0.047) and between pre-SMAm and SMAp (*p* = 0.005). Other pairwise comparisons were non-significant.

In the 50–100 ms time window, the Friedman test revealed a significant effect of stimulation target on GMFP (χ*²*(5) = 31.8, *p* < 0.001) (Figure 2B) with a moderate effect size (Kendall’s *W* = 0.303). Post-hoc analysis showed a significant difference between the most anterior (Pre-SMAa) and the most posterior (SMAp) target (*p* = 0.030). Other pairwise comparisons were non-significant (*p* > 0.05). There were no significant differences in GMFP in the 100–200 ms window (χ²(5) = 9.76, *p* = 0.082) (Figure 2C).

To quantify local reactivity in the first 50 ms, we compared the peak-to-peak amplitude between targets over the channel with the highest peak-to-peak amplitude inside the ROI. The effect of targets was significant (χ²(5) = 49.4, p < 0.001), with a moderate effect size (Kendall’s *W* = 0.471). Anterior pre-SMA exhibited significantly lower peak-to-peak values than any other target (*p* = 0.002 with pre-SMAm, *p* < 0.001 with pre-SMAp, *p* < 0.001 with pre-SMA/SMA border, *p* < 0.001 with SMAa, *p* < 0.001 with SMAp). Middle pre-SMA showed lower peak-to-peak values than preSMAp (*p* = 0.043), pre-SMA/SMA border (*p* = 0.024) and SMAp (*p* = 0.002). Peak-to-peak also differed between 50 and 100 ms after the pulse (χ²(5) = 46.0, *p* < .001; Kendall’s *W* = 0.440); post-hoc tests showed that peak-to-peak amplitude in SMAp differed significantly from pre-SMAa (*p* < 0.001), pre-SMAm (*p* < 0.001), pre-SMAp (*p* = 0.003), and the pre-SMA/SMA border (*p* = 0.021); in addition, significant differences were observed between SMAa and pre-SMAa (*p* = 0.013), pre-SMAm (*p* = 0.004), and pre-SMAp (*p* = 0.043). Finally, the pre-SMA/SMA border differed significantly from pre-SMAa (*p* = 0.028) and pre-SMAm (*p* = 0.037). No peak-to-peak differences were found in the 100–200 time window (χ²(5) = 3.69, *p* = 0.594).

As an additional proxy for local reactivity, we compared the differences within a ROI under the coil (F1, Fz, FZ1, FCz, C1, CZ). In the early time window (10–50 ms), LMFP values varied across conditions (χ²(5) = 29.8, p < 0.001), but the effect size was small (Kendall’s *W* = 0.283), and no post-hoc comparison reached significance except for the most anterior and posterior targets (pre-SMAa vs. SMAp; p = 0.027). In the 50–100 time window, LMFP was different across targets (χ²(5) = 12.4, *p* = 0.030), with small effect size (Kendall’s *W* = 0.118) and no significant post-hoc differences. LMFP did not differ across targets in the 100–200 ms time window (χ²(5) = 2.69, *p* = 0.748).

### 3.2 Spectral power and natural frequency

Distribution of the TMS-evoked frequency power varied significantly across targets. The results of the time–frequency analysis are summarized in Figure 3.

There was a significant effect of the target on the relative α power (χ²(5) = 37.54, *p* <0.001) with a moderate effect size (Kendall’s *W* = 0.358). Post-hoc pairwise comparisons indicated higher values in the most anterior target (Pre-SMAa) compared to each other target (*p* = 0.049 with Pre-SMAm, *p* < 0.001 with pre-SMAp, *p* < 0.001 with Pre-SMA/SMA border, *p* < 0.001 with SMAa and *p* < 0.01 with SMAp). Other pairwise comparisons were non-significant.

The Friedman test showed a significant effect of target on β*1* power (χ²(5) = 23.42, *p* < 0.001), with a small-to-moderate effect size (Kendall’s *W* = 0.223). Post-hoc analyses revealed higher β*1* power in SMAp compared with Pre-SMAm (*p* = 0.010) and pre-SMA/SMA border (*p* = 0.010). No other pairwise comparisons were significant.

A significant effect of target was found in the β*2* power (χ²(5) = 38.88, *p* < 0.001) with a moderate effect size (Kendall’s *W* = 0.370). Post-hoc tests showed significantly lower β*2* in pre-SMAa compared with the other pre-SMA targets (*p* <0.001 with pre-SMAm and pre-SMAp), Pre-SMA/SMA border (*p* < 0.001), and SMAa (*p* < 0.001). Also pre-SMAm showed lower β*2* levels than SMAa (*p* = 0.028).

In the γ band, the statistical test showed a significant effect of target condition (χ²(5) = 20.86, *p* = 0.023) with a small effect size (Kendall’s *W* = 0.199). Post-hoc tests revealed only one marginally significant difference between pre-SMAa and SMAa, with the first one showing higher γ power (*p* = 0.049).

Overall, TMS-evoked α power peaked over more frontal targets, whereas β*2* was predominant at the border between pre-SMA and SMA and SMAa. SMAp was instead closely linked to the β*1* rhythm. Minor effects were observed for γ, indicating a trend toward higher power over more frontal targets. Interestingly, natural frequency did not differ between targets (χ*²*(5) = 3.88, *p* = 0.567).

### 3.3 Whole-thalamic connectivity

Complementing the TMS–EEG findings, DWI analyses revealed distinct structural connectivity profiles along the SMA–pre-SMA axis (χ²(5) = 18.2, *p* < 0.001; Kendall’s *W* = 0.435) (Figure 4E). Whole-thalamic connectivity measures derived from AFD indicated significant heterogeneity across cortical targets, with pre-SMAa exhibiting lower connectivity compared to pre-SMAp (*p* = 0.002), pre-SMA/SMA border (*p* = 0.006), SMAa (*p* < 0.001), and SMAp (*p* < 0.001). SMAp consistently showed the highest connectivity, also compared with the other pre-SMA targets (*p* < 0.001 with pre-SMAm and *p* = 0.004 with pre-SMAp), suggesting stronger thalamic projections to posterior SMA regions.

**Figure 4.**
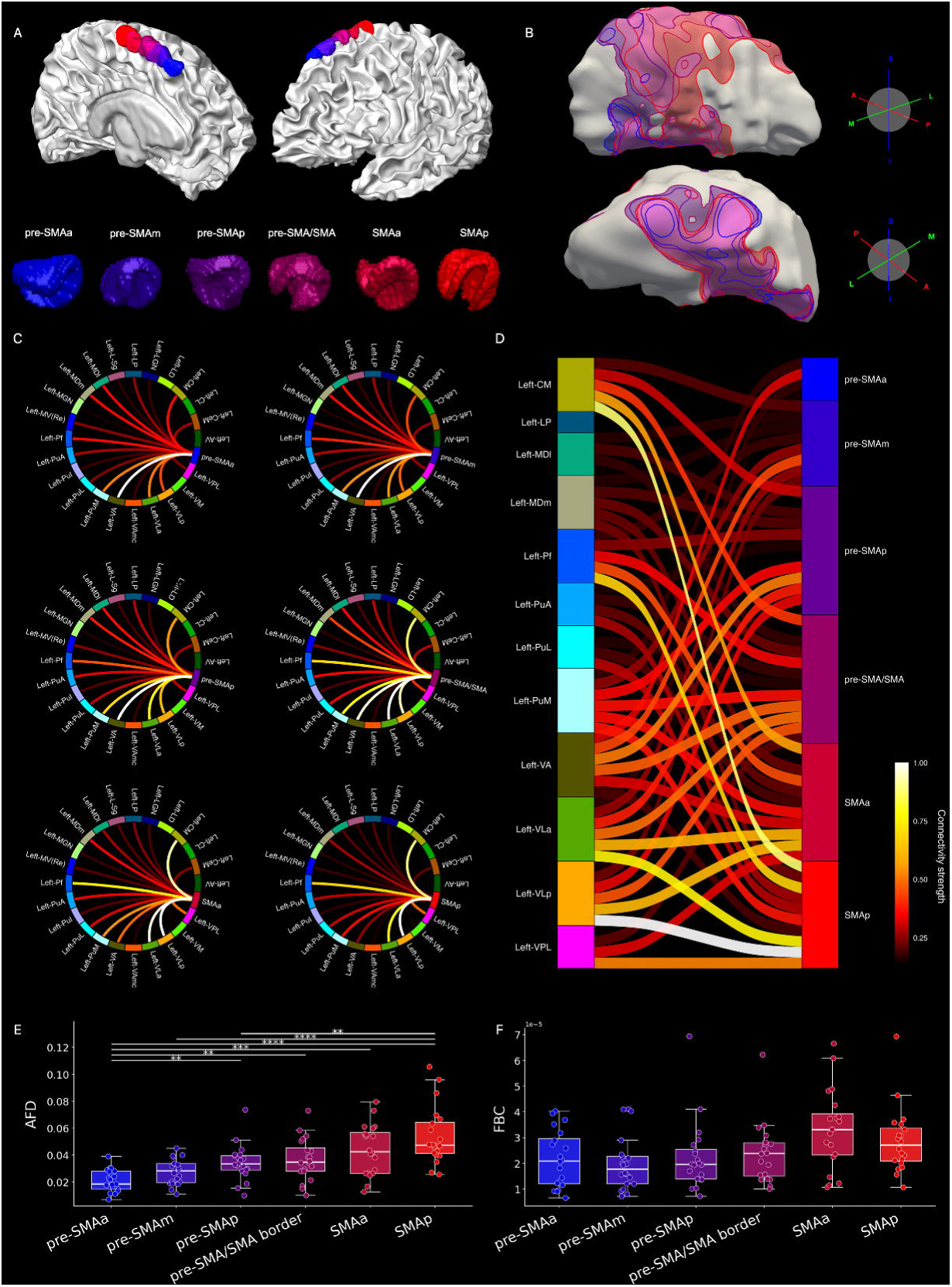
Thalamic connectivity across stimulation targets. A) Spherical ROIs with 6-mm radius localized on the white matter (WM) surface (upper part), and their individual shapes (lower part). Spheres are centered at coordinates corresponding to the maximum E-field intensity (spherical head model). B) Smoothed thalamic surface with ROI to thalamus bundles projections (A - anterior, P - posterior, M - medial, L - lateral, S - superior, I - inferior). C) Connectogram visualizations for each stimulation target, showing distinct thalamic connection patterns as the stimulation site progressed from anterior to posterior cortical locations. For each target, connectivity strengths are normalized independently (min-max scaling within that target) to highlight the relative distribution of connections. D) Sankey plot showing all stimulation targets together, with connectivity strengths normalized to the maximum value across all targets. This reveals thatanterior sites exhibited concentrated connectivity to single dominant thalamic nuclei, whereas posterior targets demonstrated broader and stronger connectivity across multiple thalamic regions. Values are normalized to the min-max value across all targets. Pre-SMAa = anterior pre-SMA, pre-SMAm = middle pre-SMA, pre-SMAp = posterior pre-SMA, SMAa = anterior SMA, SMAp = posterior SMA. E) Box plots of apparent fiber density (AFD) - derived connectivity across targets, with overlaid individual data points. F) Box plots of fiber bundle capacity (FBC) across targets, with overlaid individual data points. Color coding from blue to red indicates the anterior–posterior position of stimulation targets (blue = most anterior, red = most posterior). AV = Anteroventral nucleus; CeM = Centromedial nucleus; CL = Centrolateral nucleus; CM = Centromedian nucleus; LD = Lateral dorsal nucleus; LGN = Lateral geniculate nucleus; LP = Lateral posterior nucleus; L-Sg = Limitans-Sg nucleus; MDl, Mediodorsal lateral nucleus; MDm = Mediodorsal medial nucleus; MGN = Medial geniculate nucleus; MV(Re) = Mammillothalamic/Reuniens nucleus; Pf = Parafascicular nucleus; PuA = Pulvinar anterior nucleus; PuI = Pulvinar inferior nucleus; PuL = Pulvinar lateral nucleus; PuM = Pulvinar medial nucleus; VA = Ventral anterior nucleus; VAmc = Ventral anterior magnocellular nucleus; VLa = Ventral lateral anterior nucleus; VLp = Ventral lateral posterior nucleus; VM = Ventromedial nucleus; VPL = Ventral posterolateral nucleus.

Using a similar statistical approach, no significant differences in FBC across stimulation targets were detected when considering the whole thalamus as a single structure (Figure 4E). These findings suggest that global thalamic connectivity remains relatively stable across targets, motivating a more fine-grained analysis of FBC at the level of individual thalamic nuclei to detect region-specific differences.

Of note, considering total streamline count for the left hemisphere, we found a significant effect of target (χ²(5) = 15.1, *p* = 0.010) with a small effect size (Kendall’s *W* = 0.151). Only one post-hoc test was significant, indicating higher streamline count pre-SMAm compared with SMAa (*p* = 0.036).

### 3.4 Single-nuclei connectivity

Connectivity matrices for each stimulation target revealed different patterns of thalamic connections as the stimulation site progressed from anterior to posterior cortical locations (Figure 4C). The connectogram visualizations demonstrated distinct connectivity patterns for each target, with anterior sites showing concentrated connectivity to single dominant thalamic nuclei, while posterior targets exhibited increasingly distributed connections across multiple thalamic regions. When connectivity strengths were normalized to the maximum value across all targets to enable direct comparison, posterior stimulation sites demonstrated both a broader distribution of connections and substantially higher overall connectivity strength to the thalamus compared to anterior locations (Figure 4D).

Statistical analysis of these connectivity patterns revealed that sixteen of the twenty-three thalamic nuclei (69.6%) demonstrated statistically significant differences in FBC across targets (Figure 5), highlighting spatially selective thalamo-cortical organization within the SMA complex. Details about the statistical comparison are reported in the Supplementary materials (Table S2). The substantial proportion of nuclei exhibiting significant effects indicates that thalamic connectivity patterns vary systematically as a function of stimulation site location within the SMA. The wide range in effect sizes among significant nuclei highlights that different thalamic regions demonstrate varying degrees of spatial specificity in their connectivity with SMA subregions, with some nuclei showing highly selective connectivity profiles while others exhibit more moderate regional differentiation.

**Figure 5.**
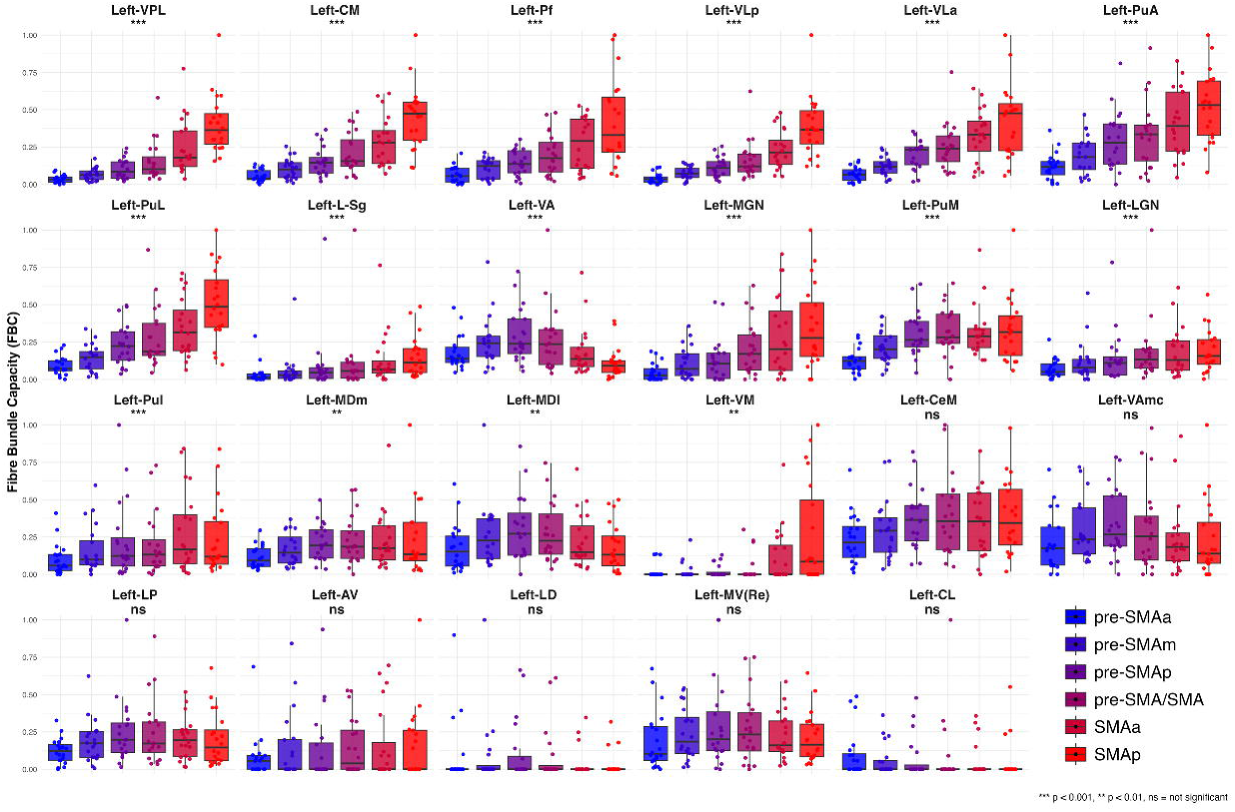
Fiber Bundle Capacity (FBC) across thalamic nuclei. Box plots show FBC values for each of the 23 thalamic nuclei. Within each nucleus panel, distributions of connectivity for each cortical target are displayed with overlaid individual data points. Target color coding from blue to red indicates anterior–posterior target position (blue = most anterior, red = most posterior). Values are normalized independently within each nucleus using min–max scaling to facilitate visualization of within-nucleus patterns. Asterisks above each boxplot indicate significant main effects of the Friedman tests across targets, with Bonferroni correction for multiple comparisons. \*\*\**p* < 0.001, \*\**p* < 0.01, ns = not significant. AV = Anteroventral nucleus; CeM = Centromedial nucleus; CL = Centrolateral nucleus; CM = Centromedian nucleus; LD = Lateral dorsal nucleus; LGN = Lateral geniculate nucleus; LP = Lateral posterior nucleus; L-Sg = Limitans-Sg nucleus; MDl, Mediodorsal lateral nucleus; MDm = Mediodorsal medial nucleus; MGN = Medial geniculate nucleus; MV(Re) = Mammillothalamic/Reuniens nucleus; Pf = Parafascicular nucleus; PuA = Pulvinar anterior nucleus; PuI = Pulvinar inferior nucleus; PuL = Pulvinar lateral nucleus; PuM = Pulvinar medial nucleus; VA = Ventral anterior nucleus; VAmc = Ventral anterior magnocellular nucleus; VLa = Ventral lateral anterior nucleus; VLp = Ventral lateral posterior nucleus; VM = Ventromedial nucleus; VPL = Ventral posterolateral nucleus.

### 3.5 Correlations between thalamo-cortical AFD-derived connectivity and GMFP

The GLS mixed-effects model revealed a strong positive association between GMFP and thalamo-target AFD-derived connectivity in the 10–50 ms time window (slope = 11.41, *p* < 0.001), indicating that regions with stronger global structural connectivity to the thalamus also exhibited heightened early cortical reactivity. This association was reduced in the 50–100 ms window, as indicated by a moderate negative change in slope relative to early time-window (Δslope = −4.36, *p* = 0.130). In contrast, in the 100–200 ms window the relationship reversed, with a large negative change in slope (Δslope = −13.85, *p* < 0.001), yielding an overall association that was comparable in magnitude but opposite in sign to that observed in the earliest window. These results show that the relationship between thalamo-target AFD-derived connectivity and TMS-evoked GMFP changes over time, reversing from positive at early latencies to negative at later latencies.

## 4. DISCUSSION

In the present study, we demonstrate with TMS–EEG and diffusion tractography that SMA is not a functionally homogeneous structure but instead exhibits different cortical excitability and oscillatory properties which are explained by distinct thalamo-cortical connectivity profiles in adjacent subregions.

Early evidence for functional heterogeneity within the SMA dates back to seminal work in the 1980s, when electrophysiological recordings and lesion studies in non-human primates suggested that adjacent regions within the medial frontal cortex could subserve partially distinct motor functions (Tanji, 1994). An anatomical distinction was hence proposed: pre-SMA as rostral portion (area F6 in non-human primates) and SMA proper as caudal portion (F3 in non-human primates; Matsuzaka et al., 1992), with their cytoarchitectonic boundary corresponding to the vertical plane intersecting the anterior commissure (Ruan et al., 2018). More recently, fMRI studies have identified distinct activation foci within the SMA and pre-SMA during resting state (Kim et al., 2010) and different motor and cognitive tasks (e.g., Tourville et al., 2019). While fMRI has been instrumental in delineating the functional topography of the medial frontal cortex, its indirect coupling to neural activity limit its ability to resolve fine-grained neurophysiological differences between closely neighboring cortical sites.

TMS-evoked global cortical excitability, as indexed by GMFP, showed clear target-dependent modulation at early latencies (10–50 ms), with posterior stimulation sites eliciting significantly stronger responses than anterior ones. Early TMS-evoked responses are often interpreted as reflecting direct activation of the stimulated cortical site (Rogasch et al., 2020). However, it is known that even within the first 50 ms post-stimulation, TMS can rapidly engage distributed cortico-cortical and subcortical circuits (Ilmoniemi et al., 1997; Komssi et al., 2002). In this regard, our findings suggest that posterior SMA regions have a greater capacity to rapidly engage distributed cortical networks following focal perturbation, which involve corticothalamic loops as key regulators of brain activity (Shine et al., 2023). We indeed found that AFD-derived connectivity with the thalamus was a strong predictor of GMFP in this early time window, in line with evidence showing network-level structural connectivity as an important contributor of TMS-evoked activity (Momi et al., 2021). Glutamatergic driver outputs from the thalamic nuclei can strongly shape cortical responses (Brandalise et al., 2025), and thalamocortical inputs have been shown to elicit stronger cortical-evoked activity than cortico-cortical inputs (Zhang & Bruno, 2019). Of note, these GMFP differences seem not attributable to global connectivity differences in the left hemisphere, since the only significant difference was pre-SMAm actually showing higher total streamline count than SMAa.

GMFP differences progressively diminished in the 50–100 ms window and were no longer present beyond 100 ms. This temporal profile suggests that the spatial specificity of TMS effects is maximal at early latencies, whereas later responses likely reflect more global, state-dependent network dynamics that are less sensitive to the specific stimulation site. During the late components (100–200 ms), thalamic contribution remained significant but exhibited much lower sensitivity, reflected in a markedly reduced slope of the linear relationship. This pattern aligns with the findings of Russo et al. (2025), who demonstrated that late evoked components critically depend on thalamocortical recurrent dynamics. Together, these observations suggest that the thalamus may play a dual role: acting as a rapid driver during the initial phase of cortical activation, while modulating distributed network dynamics in later phases. The differential sensitivity across temporal windows likely reflects the distinct underlying mechanisms, with early responses dominated by direct thalamic input and later responses shaped by network feedback and intracortical processing.

While robust target-dependent differences were observed in GMFP within the 10–50 ms window, corresponding differences in local reactivity (LMFP) were markedly smaller. This dissociation suggests that the observed early effects cannot be fully explained by differences in local cortical excitability alone, rather pointing to early, target-specific engagement of large-scale networks.

Time–frequency analyses revealed a systematic organization of oscillatory responses along the antero–posterior SMA axis. Specifically, relative α power was maximal over the most anterior pre-SMA target, consistent with the involvement of more frontal regions in higher-order cognitive control and top–down processing (Obeso et al., 2013; Palva & Palva, 2011), with an important role of GABAergic neurotransmission (Quetscher et al., 2015). The higher α may also have a structural correspondence in the higher cell density in layer V in pre-SMA (Ruan et al., 2018), pyramidal cells in neocortical layer V being an important generator of intrinsic α oscillations (Lopes da Silva, 1991). In contrast, β-band activity showed clear spatial differentiation: β2 power peaked at the intersection between pre-SMA and SMA, while β1 power was enhanced in posterior SMA. This pattern may partly reflect the closer functional coupling of SMA proper with the motor network, as β oscillations are strongly associated with motor processing. Consistent with this interpretation, SMA targets showed stronger connectivity with motor-related thalamic nuclei (VLa and VLp; Figure 5), which have causally been linked to the EEG β rhythm (Van Der Werf et al., 2006). Interestingly, natural frequency did not differ across targets, showing that this index is less sensitive to fine-grained differences in target location, probably reflecting functional properties of large-scale regions (Rosanova et al., 2009).

Functional and anatomical studies have demonstrated that pre-SMA and SMA show distinct corticocortical and cortico-subcortical connectivity profiles (Kim et al. 2010). Pre-SMA exhibits stronger connectivity with inferior frontal and posterior parietal cortices, whereas SMA proper shows stronger co-activation with the precentral gyrus and caudal dorsal premotor cortex (Ruan et al., 2018). They project to different portions of the basal ganglia, specifically of the striatum, with SMA mainly connected to the putamen and pre-SMA to the caudate (Lehericy et al., 2004; Johansen-Berg et al., 2004). These striatal territories, in turn, relay information to distinct thalamic nuclei (Postuma et al., 2006), forming partially segregated cortico–basal ganglia–thalamocortical loops. Extending this framework, our findings reveal robust connectivity between SMA proper and posterior thalamic nuclei, underscoring a critical subcortical dimension of SMA organization. The convergence of cortical motor inputs, striatal pathways conveying basal ganglia output, and distributed thalamic projections positions SMA proper as an integrative hub in which cortical and subcortical signals are combined to support coordinated voluntary movement.

A critical distinction between our analytical approaches lies in the nature of the connectivity metrics employed. While AFD-derived connectivity quantifies intra-axonal fiber volume distributed throughout individual voxels along white matter pathways (Raffelt et al., 2017), FBC provides a fundamentally different measure by quantifying the aggregate intra-axonal cross-sectional area specifically at pathway endpoints (Smith et al., 2022). This endpoint-focused measurement offers a proportional estimate of the total number of axons terminating in a given region, thereby providing a more direct assessment of the anatomical substrate underlying the capacity for information transfer between structurally distinct brain regions (Parsons et al., 2022; Sommer et al., 2017).

Our demonstration of progressively increasing AFD connectivity from pre-SMAa to SMAp, combined with the observation that 70% of thalamic nuclei exhibited statistically significant FBC differences across cortical targets, supports a hierarchical organization of thalamo-SMA interactions. Within this framework, anterior SMA subdivisions engaged more selective thalamic territories, whereas posterior regions demonstrated broader, multi-nuclear connectivity. This organization is consistent with established functional models that position pre-SMA as a cognitive–motor interface involved in action planning, decision-making, and response selection, and SMA proper as a hub more directly implicated in motor execution and sequencing (Vergani et al., 2014). Moreover, these findings align with prior evidence suggesting that human thalamic connectivity is organized into medial-to-lateral functional bands, rather than uniform nuclear units, with connectivity-defined regions spanning multiple histological nuclei to form distributed cortical networks (O’Muircheartaigh et al., 2015).

Our initial analysis, treating the thalamus as a single, unified structure, revealed no significant differences in FBC across the six stimulation targets (Figure 4E), suggesting that the overall magnitude of corticothalamic connectivity remains relatively stable regardless of the specific cortical stimulation site. However, this whole-thalamus perspective masked substantial heterogeneity in connectivity patterns. When we subsequently examined FBC at the level of individual thalamic nuclei, each nucleus demonstrated significant differences between the six stimulation targets, revealing distinct region-specific connectivity profiles that were obscured in the aggregate analysis.

These contrasting findings highlight an important principle: while global thalamic connectivity appears relatively uniform across different cortical targets, the underlying anatomical organization exhibits marked specificity at the nuclei level (Kumar et al., 2023).

Among the nuclei demonstrating significant effects, the ventral anterior (VA), ventral lateral anterior (VLa), centromedian (CM), parafascicular (PF), pulvinar medial (PuM), and mediodorsal (MD) nuclei showed particularly robust connectivity with the pre-SMA/SMA complex. The VA and VLa nuclei, embedded within basal ganglia–thalamo–cortical loops, relay basal ganglia output to premotor and supplementary motor regions to support motor preparation, sequencing, and initiation (D’Cruz et al., 2021; Petersen et al., 2023), while the intralaminar nuclei (CM/PF) play a modulatory role in arousal, salience processing, and action selection through widespread cortical and striatal projections (Zeng et al., 2024). The PuM, classically associated with attentional control and sensory integration (Soulier et al., 2023), contributes to the coordination of perceptual and motor processes. Complementing these motor and attentional systems, the MD nucleus, reciprocally interconnected with the prefrontal cortex, serves as a critical partner in executive functions including planning, cognitive control, working memory, decision-making, and cognitive flexibility (Parnaudeau et al., 2018; Wolff & Halassa, 2024). The convergent engagement of these functionally distinct thalamic nuclei with the SMA/pre-SMA complex underscores its role as an integrative cortical hub that combines motor execution, attentional control, salience-related processing, and higher-order executive functions from distributed thalamic systems (Casula et al., 2022; Guidali et al., 2023).

Our novel finding that small variations in stimulation sites can lead to substantially different neurophysiological effects may have important implications for non-invasive brain stimulation in clinical contexts, by providing a neurophysiological rationale for identifying optimal stimulation targets to engage specific networks and modulate symptoms (Lioumis et al., 2025). Different methods, based on resting state EEG, structural MRI or fMRI, have been employed to individualize targeting with the aim to maximize therapeutic benefits (see Cash & Zalesky, 2024, for a review). However, to date, there is no conclusive evidence that such personalized targeting strategies consistently outperform fixed, anatomy-based protocols derived from standard anatomical landmarks (Terao & Kodama, 2025). In this context, little attention has been paid to TMS–EEG for target optimization, likely due to its technical challenges and its sensitivity to artifacts and user performance (Lioumis & Rosanova 2022). Nonetheless, TMS–EEG offers the advantage of being a direct measure of cortical excitability, enabling the exploration of effective connectivity between nodes within the network of interest (Bortoletto et al., 2015).

Importantly, the nucleus-specific thalamo-cortical connectivity patterns identified in the present study provide a paradigm shift for therapeutic neuromodulation, suggesting that stimulation sites could be selected not only on anatomical or functional grounds, but also based on their capacity to engage specific subcortical circuits that shape network-level activity and potentially modify clinical outcomes.

Some limitations of the present study should be acknowledged. First, later sensory components of the evoked activity (Nikouline et al., 1999) may partially have contributed to some of the observed spectral differences across targets. Crucially, however, the main neurophysiological findings of the present study are based on early reactivity responses (e.g., 10–50 ms), which are temporally dissociable from late sensory-evoked components and are widely regarded as reflecting direct cortical reactivity to TMS (Rogasch et al., 2020; Tremblay et al., 2019). Therefore, non-specific responses cannot account for the robust early GMFP differences observed across stimulation sites, supporting the notion that the reported early effects reflect genuine target-specific cortical excitability rather than confounding late sensory processes.

A further limitation concerns the potential impact of EEG preprocessing on TMS-evoked responses. As with any TMS–EEG study, preprocessing steps such as artifact rejection and component suppression may influence the amplitude and spectral characteristics of the evoked signals. In the present dataset, however, data quality was generally high and only limited preprocessing was required, due to the quality control performed prior to recording (Lioumis&Rosanova, 2022; Ukharova et al., 2025). Although in a subset of participants the most anterior stimulation site exhibited a higher incidence of artifacts over frontal electrodes, necessitating the application of the SSP–SIR algorithm, this factor is unlikely to explain the systematic gradient of cortical reactivity observed across all six stimulation targets. Importantly, the observed GMFP differences were expressed at a global scalp level rather than being confined to local channels surrounding the stimulation site, further arguing against a preprocessing-driven or locally induced artifact explanation. Moreover, the consistency of the reactivity gradient across multiple targets and subjects suggests that the reported effects reflect genuine neurophysiological differences rather than differential artifact suppression.

## 5. CONCLUSION

This study demonstrates that TMS–EEG can resolve spatial differences in cortical excitability and oscillatory properties not only at the macroscale between distinct cortical areas (Rosanova et al., 2009; Passera et al., 2025) but also at a much finer spatial scale within a single functional region. The observed differences in TMS-evoked reactivity are most likely attributable to variations in connectivity patterns and cytoarchitecture of the stimulated cortical targets, rather than to differences in the induced electric field, which was kept constant across targets (see Supplementary Material for details). These findings have important clinical implications, opening new opportunities for the individualization of therapeutic interventions.

## ACKNOWLEDGEMENTS

This work was supported by Wellcome Leap as part of the Multi-Channel Psych Program. This project has also received funding from the European Research Council (ERC) under the European Union’s Horizon 2020 Research and Innovation Programme (grant agreement No 81037). Additionally, funding has been received from the Swedish Cultural Foundation and the Jane and Aatos Erkko Foundation. JG has received a personal grant from the Emil Aaltonen Foundation and a Young Group Leaders Grant from the Sigrid Juselius Foundation. DBA received funding from the Research Council of Finland (decisions 346831 and 353798). NM has received R01MH112748, R01AG042512, K24MH116366, R01MH132610, R01MH125860, R21NS136960, R01NS125307.

## DECLARATION OF INTERESTS

PL is a consultant to Nexstim Plc for TMS–EEG applications and speech cortical mapping.

RJI has patents on TMS technology, has consulted Nexstim Plc on TMS, and is co-founder and shareholder of Cortisys Ltd.

The other authors report no potential conflicts of interest.

