## Supplementary materials for "Distinct cortical excitability and connectivity profiles within the human SMA complex"

*authors corresponding equally

1. **TMS parameters**

There were no significant differences in the induced E-field intensity across targets (*χ²*(5) = 2.17, *p* = 0.825, Kendall’s *W* = 0.021) (see Supplementary Table 1). Significant differences across targets were found in stimulation depth (χ²(5) = 36.5, *p* < 0.001, Kendall’s *W* = 0.348), with pre-SMAa showing significantly lower depth than SMAa and SMAp (*p* = 0.011 for both), while all other pairwise comparisons were not significant. Also stimulation intensity, defined as the percentage of stimulation output of the TMS device, varied between targets (χ²(5) = 52.6, *p* < 0.001, Kendall’s *W* = 0.501). Post-hoc Wilcoxon signed-rank tests with Bonferroni correction indicated that pre-SMAa differed from pre-SMAp (*p* = 0.003), pre-SMA/SMA border (p = 0.003), SMAa (*p* = 0.001), and SMAp (*p* = 0.003), while pre-SMAm differed from pre-SMAp (p = 0.024), pre-SMA/SMA border (*p* = 0.017), SMAa (p = 0.009), and SMAp (p = 0.013). All other pairwise comparisons were not significant (*p* > 0.05), indicating that intensity varies between several targets, with pre-SMAa and pre-SMAm generally higher than other targets.

**Table S1. Mean TMS parameters for each stimulation target**

|  | Pre-SMAa | Pre-SMAm | Pre-SMAp | Pre-SMA/SMA border | SMAa | SMAp |
| --- | --- | --- | --- | --- | --- | --- |
| Stimulation depth (mm) | 19.96±1.66 | 20.17±1.60 | 20.20±1.57 | 20.23±1.55 | 20.35±1.62 | 20.43±1.70 |
| E-field maximum (V/m) | 91.61±10.80 | 91.87±11.37 | 92.14±10.13 | 91.68±9.93 | 92.10±9.84 | 92.16±9.95 |
| Stimulation intensity  (% MSO) | 60.33±5.55 | 59.43±6.05 | 56.71±5.63 | 56.24±5.37 | 55.62±5.81 | 55.81±6.65 |

Pre-SMAa = anterior pre-SMA, pre-SMAm = middle pre-SMA, pre-SMAp = posterior pre-SMA, SMAa = anterior SMA, SMAp = posterior SMA, MSO = maximum stimulator's output.

**Table S2. Results of Friedman tests and post-hoc pairwise comparisons assessing connectivity between pre-SMA and SMA targets and thalamic nuclei**

| **Nuclei** | **Friedman results for each nucleus** | | | | **Post-hoc pairwise comparison results** | | | |
| --- | --- | --- | --- | --- | --- | --- | --- | --- |
|  | *χ²* | Kendall’s *W* | *p* | adj-*p* | Target 1 | Target 2 | *p* | adj-*p* |
| **Left-AV** | 49.086 | 0.491 | 0.000 | 0.000*** | - | - | - | - |
| **Left-VLa** | 66.400 | 664.000 | 0.000 | 0.000*** | - | - | - | - |
|  | | | | | pre-SMAa | pre-SMA/SMA | 0.000 | 0.001** |
|  |  |  |  |  | pre-SMAa | SMAa | 0.000 | 0.001** |
|  |  |  |  |  | pre-SMAa | SMAp | 0.000 | 0.001** |
|  |  |  |  |  | pre-SMAm | SMAa | 0.000 | 0.001** |
|  |  |  |  |  | pre-SMAa | pre-SMAp | 0.000 | 0.002** |
|  |  |  |  |  | pre-SMAm | SMAp | 0.000 | 0.002** |
|  |  |  |  |  | pre-SMAa | pre-SMAm | 0.000 | 0.005** |
|  |  |  |  |  | pre-SMAm | pre-SMAp | 0.000 | 0.006** |
|  |  |  |  |  | pre-SMAp | SMAp | 0.001 | 0.008** |
|  |  |  |  |  | pre-SMAp | SMAa | 0.001 | 0.010** |
|  |  |  |  |  | pre-SMAm | pre-SMA/SMA | 0.001 | 0.014* |
| **Left-PuM** | 34.000 | 0.340 | 0.000 | 0.000*** | - | - | - | - |
|  | | | | | pre-SMAa | SMAa | 0.000 | 0.002** |
|  |  |  |  |  | pre-SMAa | pre-SMAp | 0.000 | 0.002** |
|  |  |  |  |  | pre-SMAa | pre-SMA/SMA | 0.000 | 0.004** |
|  |  |  |  |  | pre-SMAa | pre-SMAm | 0.000 | 0.006** |
|  |  |  |  |  | pre-SMAa | SMAp | 0.001 | 0.019* |
|  |  |  |  |  | pre-SMAm | pre-SMAp | 0.002 | 0.035* |
| **Left-PuA** | 61.657 | 0.617 | 0.000 | 0.000*** | - | - | - | - |
|  | | | | | pre-SMAa | SMAa | 0.000 | 0.001** |
|  |  |  |  |  | pre-SMAa | SMAp | 0.000 | 0.001** |
|  |  |  |  |  | pre-SMAm | SMAp | 0.000 | 0.001** |
|  |  |  |  |  | pre-SMAm | SMAa | 0.000 | 0.002** |
|  |  |  |  |  | pre-SMAa | pre-SMAp | 0.000 | 0.004** |
|  |  |  |  |  | pre-SMAa | pre-SMA/SMA | 0.000 | 0.005** |
|  |  |  |  |  | pre-SMAa | pre-SMAm | 0.001 | 0.021* |
|  |  |  |  |  | pre-SMAm | pre-SMAp | 0.001 | 0.024* |
|  |  |  |  |  | pre-SMAp | SMAp | 0.002 | 0.045* |
| **Left-VPL** | 76.743 | 0.767 | 0.000 | 0.000* | - | - | - | - |
|  | | | | | pre-SMAa | pre-SMA/SMA | 0.000 | 0.001** |
|  |  |  |  |  | pre-SMAa | SMAa | 0.000 | 0.001** |
|  |  |  |  |  | pre-SMAa | SMAp | 0.000 | 0.001** |
|  |  |  |  |  | pre-SMAm | SMAa | 0.000 | 0.001** |
|  |  |  |  |  | pre-SMAm | SMAp | 0.000 | 0.001** |
|  |  |  |  |  | pre-SMAp | SMAa | 0.000 | 0.001** |
|  |  |  |  |  | pre-SMAp | SMAp | 0.000 | 0.001** |
|  |  |  |  |  | pre-SMA/SMA | SMAp | 0.000 | 0.001** |
|  |  |  |  |  | pre-SMAa | pre-SMAp | 0.001 | 0.002** |
|  |  |  |  |  | pre-SMAa | pre-SMAm | 0.001 | 0.004** |
|  |  |  |  |  | pre-SMA/SMA | SMAa | 0.001 | 0.006** |
|  |  |  |  |  | pre-SMAm | pre-SMAp | 0.002 | 0.008** |
|  |  |  |  |  | pre-SMAm | pre-SMA/SMA | 0.001 | 0.013* |
|  |  |  |  |  | SMAa | SMAp | 0.001 | 0.024* |
| **Left-L-Sg** | 53.351 | 0.534 | 0.000 | 0.000*** | - | - | - | - |
|  | | | | | pre-SMAa | SMAp | 0.000 | 0.001** |
|  |  |  |  |  | pre-SMAa | SMAa | 0.000 | 0.003** |
|  |  |  |  |  | pre-SMAa | pre-SMAp | 0.001 | 0.014* |
|  |  |  |  |  | pre-SMAm | SMAa | 0.001 | 0.016* |
|  |  |  |  |  | pre-SMAm | SMAp | 0.001 | 0.019* |
|  |  |  |  |  | pre-SMAa | pre-SMA/SMA | 0.002 | 0.030* |
|  |  |  |  |  | pre-SMAp | SMAp | 0.002 | 0.035* |
| **Left-LD** | 5.669 | 0.057 | 0.340 | 1.000 | - | - | - | - |
| **Left-MGN** | 41.269 | 0.413 | 0.000 | 0.000*** | - | - | - | - |
|  | | | | | pre-SMAa | SMAp | 0.000 | 0.003** |
|  |  |  |  |  | pre-SMAp | SMAp | 0.000 | 0.006** |
|  |  |  |  |  | pre-SMAa | SMAa | 0.000 | 0.007** |
|  |  |  |  |  | pre-SMAm | SMAp | 0.001 | 0.014* |
|  |  |  |  |  | pre-SMAa | pre-SMA/SMA | 0.001 | 0.018* |
|  |  |  |  |  | pre-SMAm | SMAa | 0.002 | 0.024* |
|  |  |  |  |  | pre-SMAp | SMAa | 0.002 | 0.034* |
| **Left-CeM** | 18.486 | 0.185 | 0.002 | 0.055 | - | - | - | - |
| **Left-VLp** | 76.743 | 0.767 | 0.000 | 0.000*** | - | - | - | - |
|  | | | | | pre-SMAa | SMAa | 0.000 | 0.001** |
|  |  |  |  |  | pre-SMAa | SMAp | 0.000 | 0.001** |
|  |  |  |  |  | pre-SMAm | SMAp | 0.000 | 0.001** |
|  |  |  |  |  | pre-SMAp | SMAp | 0.000 | 0.001** |
|  |  |  |  |  | pre-SMAm | SMAa | 0.000 | 0.002** |
|  |  |  |  |  | pre-SMAp | SMAa | 0.000 | 0.002** |
|  |  |  |  |  | pre-SMAa | pre-SMA/SMA | 0.000 | 0.002** |
|  |  |  |  |  | pre-SMAa | pre-SMAp | 0.000 | 0.003** |
|  |  |  |  |  | pre-SMAa | pre-SMAm | 0.000 | 0.005** |
|  |  |  |  |  | SMAa | SMAp | 0.000 | 0.005** |
|  |  |  |  |  | pre-SMA/SMA | SMAp | 0.000 | 0.006** |
|  |  |  |  |  | pre-SMAm | pre-SMAp | 0.001 | 0.019* |
|  |  |  |  |  | pre-SMAm | pre-SMA/SMA | 0.001 | 0.019* |
| **Left-VA** | 49.086 | 0.491 | 0.000 | 0.000*** | - | - | - | - |
|  | | | | | pre-SMAm | SMAp | 0.000 | 0.002** |
|  |  |  |  |  | pre-SMAp | SMAp | 0.000 | 0.002** |
|  |  |  |  |  | pre-SMA/SMA | SMAp | 0.000 | 0.002** |
|  |  |  |  |  | pre-SMAp | SMAa | 0.000 | 0.005** |
|  |  |  |  |  | SMAa | SMAp | 0.000 | 0.005** |
|  |  |  |  |  | pre-SMAa | pre-SMAp | 0.001 | 0.010** |
|  |  |  |  |  | pre-SMA/SMA | SMAa | 0.001 | 0.010** |
|  |  |  |  |  | pre-SMAa | pre-SMAm | 0.001 | 0.011* |
|  |  |  |  |  | pre-SMAm | SMAa | 0.003 | 0.040* |
| **Left-MDl** | 24.600 | 246.000 | 0.000 | 0.004** | - | - | - | - |
|  | | | | | pre-SMAa | pre-SMAp | 0.000 | 0.003** |
|  |  |  |  |  | pre-SMAa | pre-SMAm | 0.000 | 0.007** |
| **Left-LGN** | 31.094 | 0.311 | 0.000 | 0.000*** | - | - | - | - |
|  | | | | | pre-SMAa | SMAp | 0.000 | 0.006** |
|  |  |  |  |  | pre-SMAa | pre-SMA/SMA | 0.000 | 0.007** |
|  |  |  |  |  | pre-SMAa | SMAa | 0.001 | 0.016* |
|  |  |  |  |  | pre-SMAa | pre-SMAp | 0.002 | 0.024* |
| **Left-PuI** | 29.052 | 0.291 | 0.000 | 0.001*** | - | - | - | - |
|  | | | | | pre-SMAa | SMAa | 0.000 | 0.006** |
|  |  |  |  |  | pre-SMAa | pre-SMAm | 0.000 | 0.006** |
|  |  |  |  |  | pre-SMAa | SMAp | 0.000 | 0.007** |
|  |  |  |  |  | pre-SMAa | pre-SMAp | 0.001 | 0.019* |
| **Left-VM** | 23.448 | 0.234 | 0.000 | 0.006** | - | - | - | - |
| **Left-MV(Re)** | 9.770 | 0.098 | 0.082 | 1.000 | - | - | - | - |
| **Left-CL** | 6.288 | 0.063 | 0.279 | 1.000 | - | - | - | - |
| **Left-VAmc** | 14.347 | 0.143 | 0.014 | 0.312 | - | - | - | - |
| **Left-MDm** | 26.971 | 0.270 | 0.000 | 0.001** | - | - | - | - |
|  | | | | | pre-SMAa | pre-SMAp | 0.000 | 0.002** |
|  |  |  |  |  | pre-SMAa | pre-SMAm | 0.001 | 0.010** |
|  |  |  |  |  | pre-SMAa | pre-SMA/SMA | 0.001 | 0.013* |
|  |  |  |  |  | pre-SMAa | SMAa | 0.001 | 0.016* |
| **Left-PuL** | 59.229 | 0.592 | 0.000 | 0.001*** | - | - | - | - |
|  | | | | | pre-SMAa | pre-SMA/SMA | 0.000 | 0.001** |
|  |  |  |  |  | pre-SMAa | SMAp | 0.000 | 0.002** |
|  |  |  |  |  | pre-SMAa | pre-SMAp | 0.000 | 0.002** |
|  |  |  |  |  | pre-SMAa | SMAa | 0.000 | 0.002** |
|  |  |  |  |  | pre-SMAm | SMAa | 0.000 | 0.002** |
|  |  |  |  |  | pre-SMAm | SMAp | 0.000 | 0.002** |
|  |  |  |  |  | pre-SMAm | pre-SMAp | 0.000 | 0.007** |
|  |  |  |  |  | pre-SMAp | SMAp | 0.001 | 0.016* |
|  |  |  |  |  | pre-SMAm | pre-SMA/SMA | 0.002 | 0.027* |
| **Left-Pf** | 79.029 | 0.790 | 0.000 | 0.000*** | - | - | - | - |
|  | | | | | pre-SMAa | pre-SMA/SMA | 0.000 | 0.001** |
|  |  |  |  |  | pre-SMAa | SMAa | 0.000 | 0.001** |
|  |  |  |  |  | pre-SMAa | SMAp | 0.000 | 0.001** |
|  |  |  |  |  | pre-SMAm | SMAp | 0.000 | 0.001** |
|  |  |  |  |  | pre-SMAm | SMAa | 0.000 | 0.002** |
|  |  |  |  |  | pre-SMA/SMA | SMAp | 0.000 | 0.003** |
|  |  |  |  |  | pre-SMAm | pre-SMA/SMA | 0.000 | 0.004** |
|  |  |  |  |  | pre-SMAp | SMAp | 0.000 | 0.004** |
|  |  |  |  |  | pre-SMAp | SMAa | 0.001 | 0.008** |
|  |  |  |  |  | pre-SMAm | pre-SMAp | 0.001 | 0.010** |
|  |  |  |  |  | pre-SMAa | pre-SMAm | 0.001 | 0.016* |
|  |  |  |  |  | SMAa | SMAp | 0.002 | 0.024* |
| **Left-CM** | 82.086 | 0.821 | 0.000 | 0.000*** | - | - | - | - |
|  | | | | | pre-SMAa | pre-SMA/SMA | 0.000 | 0.001** |
|  |  |  |  |  | pre-SMAa | SMAa | 0.000 | 0.001** |
|  |  |  |  |  | pre-SMAa | SMAp | 0.000 | 0.001** |
|  |  |  |  |  | pre-SMAm | SMAa | 0.000 | 0.001** |
|  |  |  |  |  | pre-SMAm | SMAp | 0.000 | 0.001** |
|  |  |  |  |  | pre-SMAp | SMAa | 0.000 | 0.00** |
|  |  |  |  |  | pre-SMAp | SMAp | 0.000 | 0.002** |
|  |  |  |  |  | pre-SMAa | pre-SMAp | 0.000 | 0.002** |
|  |  |  |  |  | pre-SMAm | pre-SMA/SMA | 0.000 | 0.004** |
|  |  |  |  |  | pre-SMAm | pre-SMAp | 0.000 | 0.005** |
|  |  |  |  |  | pre-SMA/SMA | SMAp | 0.000 | 0.005** |
|  |  |  |  |  | pre-SMAa | pre-SMAm | 0.000 | 0.005** |
|  |  |  |  |  | SMAa | SMAp | 0.001 | 0.019* |
| **Left-LP** | 13.571 | 0.136 | 0.019 | 0.427 | - | - | - | - |

Left panel: Friedman test results for each nucleus, including the Chi-Squared values, Kendall's *W* coefficients, and their corresponding unadjusted and adjusted p-values (Bonferroni-corrected). Right panel: Post-hoc pairwise comparisons within each nucleus and corresponding p-values. ****p* < 0.001: ***, ***p* < 0.01, **p* < 0.05. Pre-SMAa = anterior pre-SMA, pre-SMAm = middle pre-SMA, pre-SMAp = posterior pre-SMA, SMAa = anterior SMA, SMAp = posterior SMA.
